# Early events in midzone formation stabilize nascent bipolar spindles

**DOI:** 10.1101/2024.09.09.612135

**Authors:** Shannon Sim, Sean Moore, Khalid Al-Naemi, Ziad El-Hajj, Jackie Vogel

## Abstract

The formation of a stable mitotic spindle is critical to the accurate partitioning of the chromosomes during mitosis. Interpolar microtubules of the spindle midzone consist of antiparallel microtubules crosslinked by kinesin-5 and stabilize the spindle by opposing forces produced when sister chromatids are attached to microtubules and under tension. Despite the importance of the interpolar microtubules, how and when they form and what determines their number remain unknown. In this study, we report that a γ-tubulin mutation (γtub-Y445D) disrupts the localization of kinesin-5 and the formation of the interpolar microtubules, resulting in spindle instability. We find that kinesin-5 crosslinking is intact in this mutant, but that it is incapable of the subsequent kinesin-5 microtubule sliding needed to stabilize the nascent spindle. Early activation of the PRC1 homolog Ase1 restores nascent spindle stability to the γtub-Y445D mutant but cannot stabilize spindles during centromere attachment. Our work shows that midzone assembly begins with the formation of interpolar microtubule precursors in monopolar spindles that persist until early metaphase and limit the formation of kinetochore attachments in new spindles.

## INTRODUCTION

A stable bipolar mitotic spindle ensures the accurate segregation of genetic material into daughter cells at the end of cell division. Kinetochore microtubules (kMTs) emanating from opposing spindle poles must attach to each pair of sister chromatids to pull the chromosomes apart during anaphase. However, the interpolar microtubules (ipMTs) are also critical, as they form the midzone that stabilizes the spindle and promotes interactions between the kMTs and the chromosomes.

In budding yeast, spindle formation relies on pairs of anti-parallel microtubules (MTs) that are crosslinked by kinesin-5 motor proteins^1^ (Figure 1A). The structure and behavior of the newly formed “nascent” bipolar spindle is distinct from that of the metaphase spindle in two key aspects. First, the nascent spindle lacks a defined midzone^2,3^ and instead the bipolar state is maintained by a more variable number of ipMT precursors (pre-ipMTs) characterized by overlaps of ∼100nm^2,3^ that are short relative to the ∼400 nm overlaps characteristic of the ipMTs of the metaphase spindle^3,4^. Second, the inward forces produced by kinetochore attachments on the kinetochores of sister chromatids are expected to be low in nascent spindles and increase during metaphase as sister chromatid biorientation proceeds and tension is applied to the kinetochores. As the kMTs attach to and biorient sister chromatids, the inward forces increase and must be balanced by ipMTs. To maintain the bipolar state, the pre-ipMTs must be converted into ipMTs that are able to oppose these forces.

**Figure 1.**
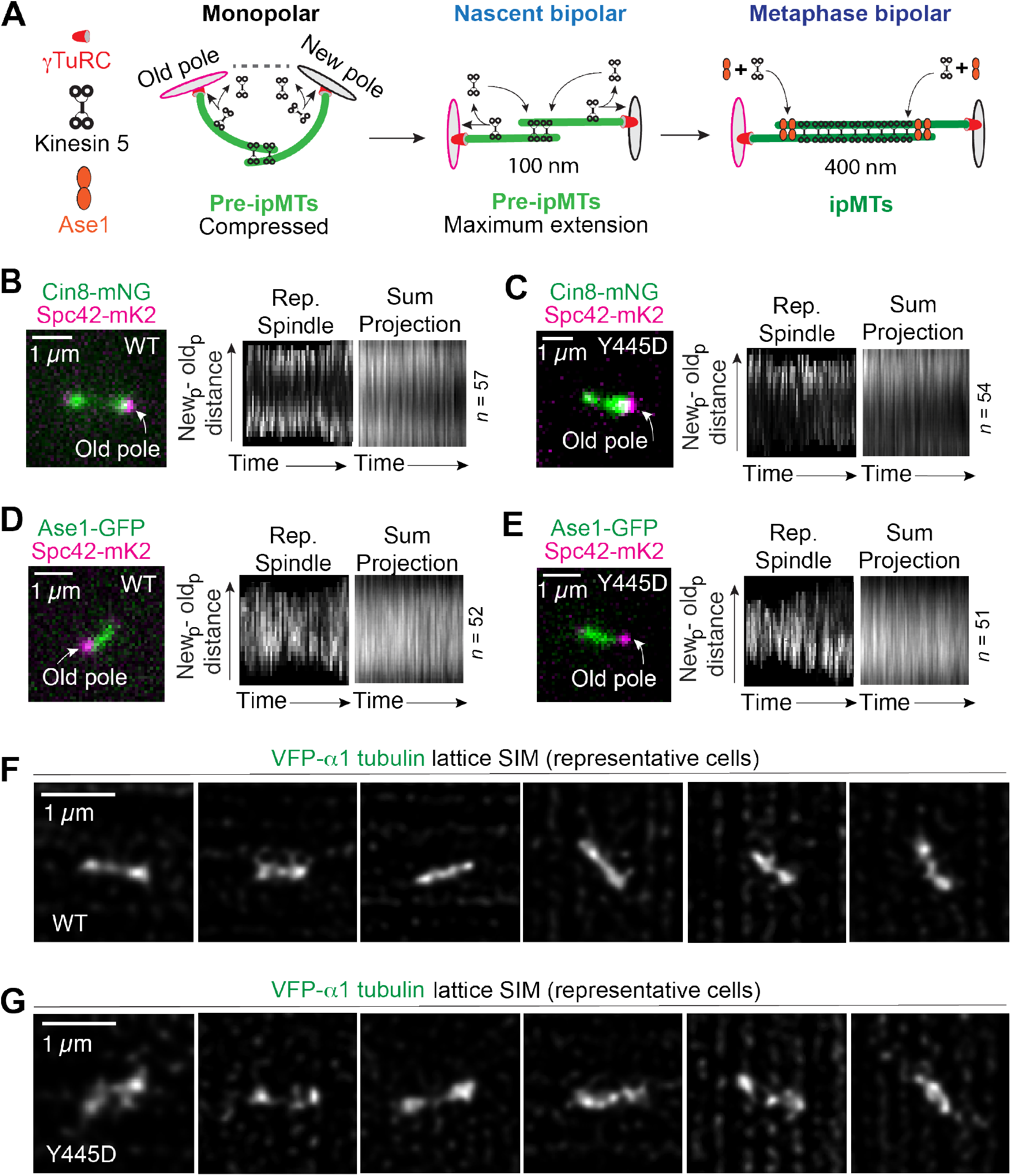
The **γ**-tubulin Y445D mutation disrupts kinesin-5 localization and ipMT formation (A) Kinesin-5 crosslinking forms pre-ipMTs in monopolar spindles that are maximally extended in nascent bipolar spindles. Recruitment of kinesin-5 and Ase1 during metaphase converts unstable pre-ipMTs into stable ipMTs. (B)-(C) Dynamic localization of Cin8-mNG during metaphase in WT and γtub-Y445D cells. Kymographs for a single representative spindle and a summed projection are shown for each condition. (D)-(E) Dynamic localization of Ase1-GFP during metaphase in in WT and γtub-Y445D cells. Kymographs for a single representative spindle and a summed projection are shown for each condition. (F)-(G) Lattice SIM analysis of midzone structure in WT and γtub-Y445D cells.

The ipMTs are essential for spindle function in metaphase and anaphase, yet how they are formed and what the requirements are for spindle function during metaphase remain largely uninterrogated. For example, it is unclear if nascent spindles can support the formation of stable kMT attachments to sister chromatids. Budding yeast offers a set of powerful tools that when used in combination allow the process of ipMT formation and the attachment of kMTs to sister chromatids to be investigated (Figure 1A). The organization and number of MTs in budding yeast are highly stereotyped and also less complex than those of metaphase spindles in animal cells^5^. The structures of both nascent (length 0.8 – 1 µm) and late metaphase (length ∼2 µm) spindles have been characterized using tomography and both the number and nanoscale organization of the pre-ipMTs and ipMTs are known for both states. The contributions of kinesin-5 to both spindle formation and stability have also been described^2,6^. Finally, precise detection of kMT attachments capable of exerting tension is possible using centromere reporters^7,8^ and super-resolution microscopy.

The minimal overlap between MTs in monopolar and nascent bipolar spindles is expected to promote the minus end-directed stepping modality described for kinesin-5 on single MTs^9–11^. As a result, the deployment of kinesin-5 to short overlaps is likely affected by activities at both the plus and minus ends of spindle MTs. This is evident in the subtle bias of kinesin-5 to MTs that assemble from the old spindle pole^12^, which would recruit kinesin-5 prior to the MTs assembled at the new spindle pole. All spindle MTs originate from γ-tubulin-Ring Complexes (γTuRCs) that are located on the inner plaque of the spindle pole body, which serves as the centrosome equivalent in fungal cells. Surprisingly, we identified mutations in γ-tubulin residues phosphorylated in vivo^13,14^ that reduce the number of ipMTs but not the total number of spindle MTs. These mutants were found to have increased rates of chromosome mis-segregation^15^ and increased spindle instability^3,6^. In particular, a phospho-mimetic mutation (Y445D) in the intrinsically disordered carboxyl terminus of budding yeast γ-tubulin (γCT)^16,17^ inhibits the breakdown of the midzone of anaphase spindles^14^ and decreases the stability of metaphase spindles^6^. Genetic evidence^6^ suggested that the γtub-Y445D mutation reduces the function of Cin8, a kinesin-5 that plays an important role in spindle formation and contributes to the crosslinking of ipMTs throughout mitosis^2,18^. Interestingly, the function of PRC1/Ase1, a non-motor MT crosslinker, was not reduced in the mutant, suggesting a specific defect in kinesin-5 during the formation of the ipMTs.

Kinesin-5 crosslinking, not sliding, is required for initial spindle formation^2^. However, sliding is required for the nascent spindle to elongate to a stable equilibrium length between 1.0 and 1.4 µm. We hypothesized that the γtub-Y445D mutant would allow us to isolate the crosslinking and sliding activities of kinesin-5 that contribute to forming the pre-ipMTs and their conversion to ipMTs, and generally gain insight into critical steps in midzone assembly that are required for initiating sister chromatid biorientation.

In this study we show that the γtub-Y445D mutation disrupts kinesin-5 localization as well as ipMT formation, and that this change is correlated with an increase in instability of both nascent and asynchronous spindles. Furthermore, we show that inhibiting Ase1 phosphorylation can maintain nascent spindle stability in the absence of kinesin-5 sliding but is unable to stabilize late metaphase spindles during centromere attachment. This suggests an inability to convert the pre-ipMTs into ipMTs through kinesin-5 sliding in this mutant.

## RESULTS

### Kinesin-5 is depleted from the spindle midzone in the phospho-mimetic γtub-Y445D mutant

Mutations in kinesin-5 are known to cause metaphase spindle instability and collapse^19–22^. Our previous work suggests that metaphase spindle instability in the γtub-Y445D mutant arises from a defect in one or both budding yeast kinesin-5 motor proteins, Cin8 and Kip1.

We used simultaneous dual-colour confocal imaging to investigate the localization of both Cin8 and Kip1 in metaphase spindles. We found that Cin8-mNeonGreen (mNG, from^23^) is biased to the old spindle pole and depleted from the spindle midzone in γtub- Y445D cells (Figure 1B and 1C). This phenotype is not observed in the phospho- inhibiting γtub-Y445F mutant (Figure S1). This confirms that the spindle instability found in the γtub-Y445D mutant correlates with a defect in Cin8 localization in pre-anaphase spindles. We found that Kip1-GFP (from^24^) is also biased to the old spindle pole and depleted from the midzone in γtub-Y445D cells (Figure S2). In contrast, we found that Ase1-GFP localization was not altered in the γtub-Y445D mutant (Figure 1D and 1E). The localization of both forms of kinesin-5 is altered in this mutant, but not that of the MT crosslinker Ase1. This suggests that the γtub-Y445D mutation specifically inhibits the recruitment of both kinesin-5 motor proteins to the MTs of the spindle midzone, while Ase1 localization is unchanged.

Cin8 phosphoregulation by Cdk1 tunes its association and dissociation from spindle MTs^25,26^. Its affinity for MTs is essential to its function in spindle formation^27^. The binding affinity of Cin8 to spindle MTs is negatively regulated by phosphorylation^25^. This is contrary to other kinesin-5s, including the human kinesin-5 Eg5, in which increased MT binding has been shown upon phosphorylation^28–31^.

We hypothesized that inhibiting Cin8 phosphorylation in the γtub-Y445D mutant would increase the bias of Cin8 to the old pole by increasing its binding to spindle MTs. We used a mutant in which phosphorylation is inhibited at the three Cdk1 phospho-sites (cin8-3A)^25^. Indeed, we found that there is an increased bias of Cin8-mNG to the old pole in cin8-3A; γtub-Y445D cells relative to both cin8-3A and γtub-Y445D cells (Figure 1C and S3). The stability of cin8-3A; γtub-Y445D pre-anaphase spindles is also not significantly different from that of γtub-Y445D spindles (Figure S3, *p* = 0.8597).

### The γtub-Y445D mutant disrupts the formation of the ipMTs

We hypothesized that the disruption in kinesin-5 localization in γtub-Y445D spindles would be corelated with a decrease in the number of ipMTs. Using lattice SIM microscopy, we visualized the structure of bipolar spindles in WT and γtub-Y445D cells (Figure 1F and 1G). We found the Tub1-venus signal to be decreased at the midzone in γtub-Y445D cells (Figure 1F and 1G). Furthermore, off-axis MTs were frequently observed in γtub-Y445D cells (Figure 1F and 1G). Slightly more Tub1-venus signal was observed at the midzone of γtub-Y445F cells (Figure S1). Thus, the disruption of kinesin-5 localization on γtub-Y445D spindles is correlated with a disruption of ipMT formation.

### The **y**tub-Y445D mutation causes spindle instability in both nascent and asynchronous bipolar spindles

We have previously shown that metaphase γtub-Y445D spindles are unstable^6^. Using a fluorophore-switching method that allows us to visualize the old and new spindle poles in different colors (Figure 2A), we measured spindle length before, during, and after the monopolar-to-bipolar transition in WT and γtub-Y445D cells.

**Figure 2.**
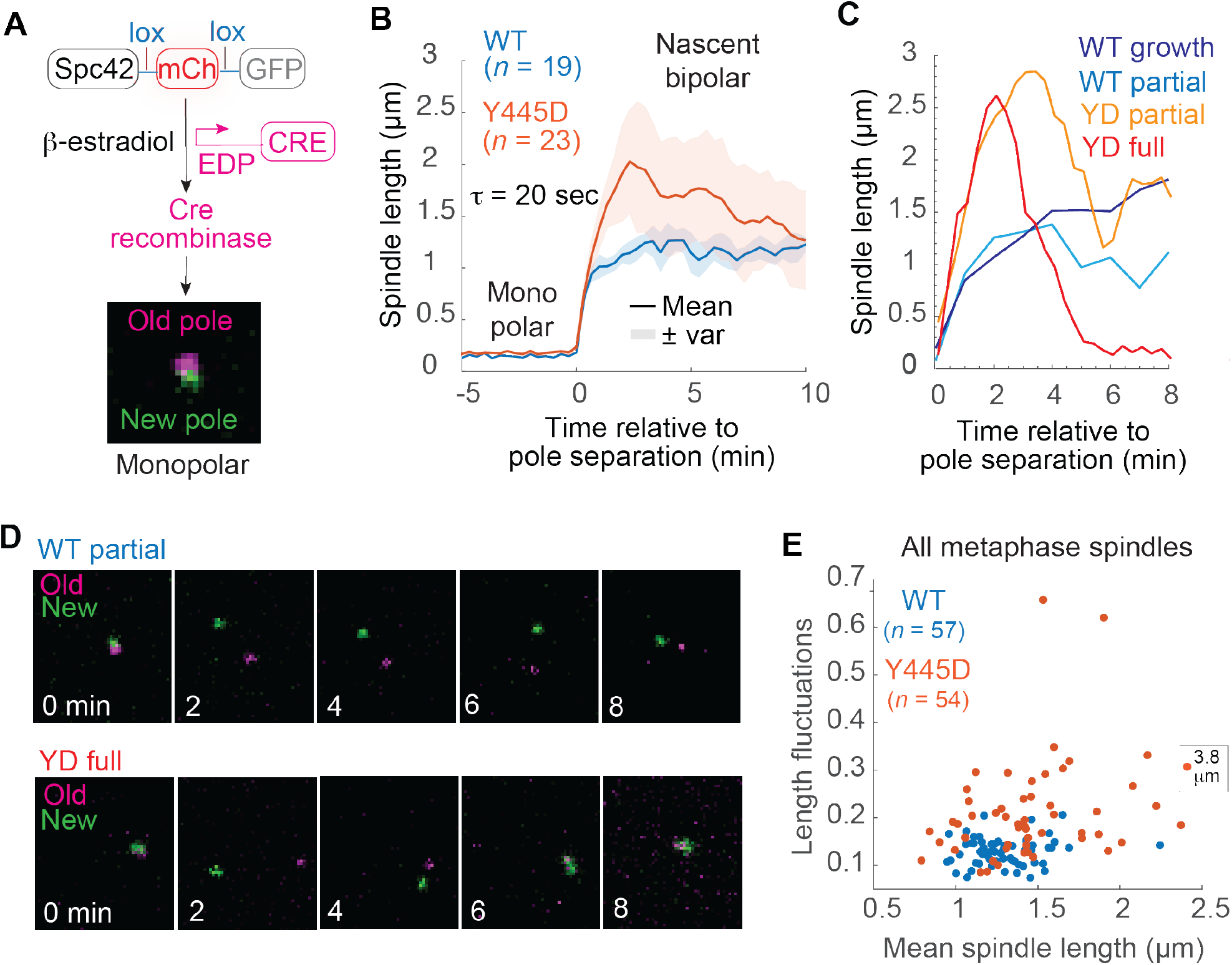
Analysis of spindle formation and stability during metaphase in **γ**tub-Y445D cells reveals a defect in converting pre-ipMTs to ipMTs (A) Method for quantitative analysis of the monopolar-to-bipolar transition using spectral separation of the old (red; mCherry) and new (green; GFP) spindle poles. (B) Average and variance of individual WT and γtub-Y445D trajectories aligned at the monopolar-to-bipolar transition. (C) Characteristic trajectories for growth and partial collapse in WT nascent spindles, as well as partial and full collapse in γtub-Y445D nascent spindles. (D) Timelapse images depicting Spc42-mKate2-mCherry and Spc42-mKate2-GFP signal for the first 8 minutes following bipolar spindle formation for a WT spindle undergoing partial collapse and a γtub-Y445D spindle undergoing full collapse. (E) Spindle length fluctuations as a function of mean spindle length for individual asynchronous WT and γtub- Y445D cells.

We found that nascent γtub-Y445D bipolar spindles were much more unstable than WT nascent spindles and collapsed more frequently (Figure 2B). Partial collapse of nascent spindles was observed in both WT and γtub-Y445D cells but was more pronounced in γtub-Y445D spindles (Figure 2C). This type of collapse was found to occur at a similar frequency in WT and γtub-Y445D spindles (Figure 2C and 2D, 38% of WT spindles and 30% of γtub-Y445D spindles). However, full collapse to a length of less than 300 nm was only observed in the γtub-Y445D mutant (Figure 2C and 2D, 26% of spindles). Furthermore, as previously reported, asynchronous γtub-Y445D bipolar spindles are significantly more unstable than WT spindles^6^ (Figure 2E, *p* = 1×10^-7^). The mis-localization of kinesin-5 and disruption of ipMT formation in the γtub-Y445D mutant are therefore correlated with increased spindle instability in both nascent and asynchronous bipolar spindles. In contrast, the stability of asynchronous γtub-Y445F spindles is consistent with that of WT spindles (Figure S1, *p* = 0.1529).

### Kinesin-5 crosslinking is intact in the **y**tub-Y445D mutant

Initial spindle pole separation is driven by kinesin-5 crosslinking of the pre-ipMTs that form short overlaps intersecting at a wide range of angles in a geometry that is not conducive to kinesin-5 sliding^2^. We investigated whether the crosslinking of the pre-ipMTs that drives the formation of the bipolar spindle was disrupted in the γtub-Y445D mutant.

We reasoned that a loss in Cin8 crosslinking function would delay or block the separation of the old and new poles (Figure 3A and 3B). We used the exclusion of the G1 transcriptional repressor Whi5 from the nucleus as a marker for commitment to enter the cell cycle (START) and an increase in the G1 Cdk1 activity required to cleave the bridge structure linking the old and new spindle poles^32–34^. However, we found that the time interval between START and bipolar spindle formation is unchanged in the γtub-Y445D mutant (Figure 3C, *p* = 0.5088), with a mean interval of 43 ± 14.2 minutes for WT cells and of 44 ± 9.2 minutes for γtub-Y445D cells.

**Figure 3.**
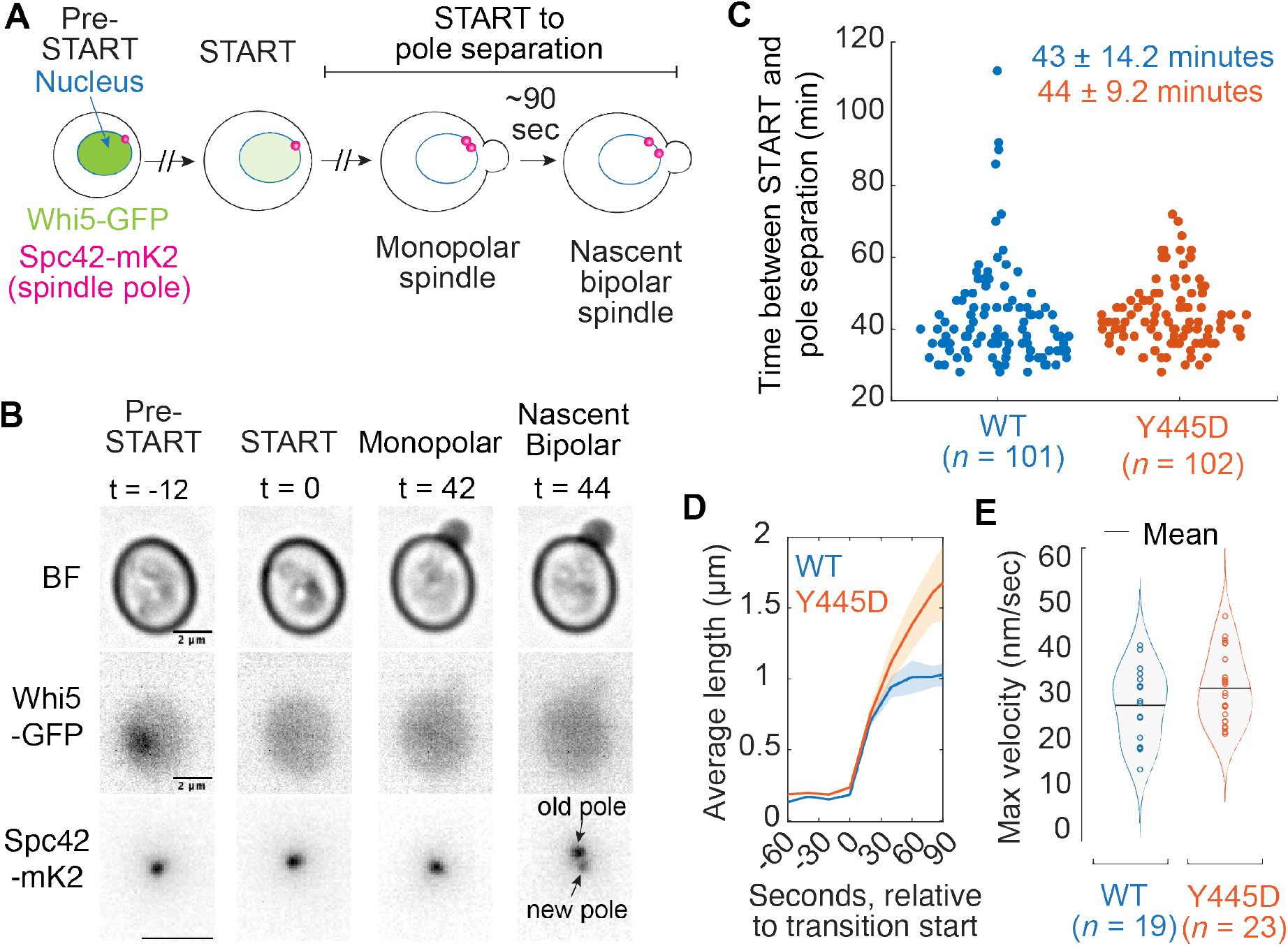
The timing of bipolar spindle formation is unchanged in **γ**tub-Y445D cells (A) Cartoon depicting how the export of Whi5-GFP from the nucleus is used to measure the time between START and bipolar spindle formation. (B) Timelapse images depicting Whi5-GFP and Spc42-mKate2 signal before and after bipolar spindle formation in a representative cell. (C) Time elapsed between START and bipolar spindle formation for individual WT and γtub-Y445D cells. (D) Average length as a function of time during the monopolar-to-bipolar transition for WT and γtub-Y445D cells. (E) Maximum velocity of spindle pole separation during the monopolar-to-bipolar transition for individual WT and γtub-Y445D cells.

We also expected that a defect in Cin8 crosslinking would decrease the pole separation velocity during the monopolar-to-bipolar transition. Using the fluorophore-switching method described previously (Figure 2A), we found the maximum velocity of the monopolar-to-bipolar transition in WT cells to be 28.6 ± 8.0 nm s^-1^, consistent with our previously published results^2^ (Figure 3D and 3E). However, we found that the maximum velocity of the transition is not significantly different in the γtub-Y445D mutant, with a value of 32.4 ± 7.6 nm s^-1^ (Figure 3D and 3E, *p* = 0.1221). Together, these results suggest that the pre-ipMTs form in the γtub-Y445D mutant.

### Early activation of Ase1 inhibits the slow growth of nascent spindles that is driven by kinesin-5 sliding

Ase1 contributes to maintaining the ipMTs, and when dephosphorylated in anaphase, it is recruited to the spindle^35,36^. Mutations that inhibit Ase1 phosphorylation (ase1-7A) are therefore expected to allow its recruitment to the pre-ipMTs of nascent bipolar spindles.

We found that the maximum velocity of the monopolar-to-bipolar transition is significantly decreased in ase1-7A cells, with a value of 21.0 ± 4.8 nm s^-1^ (Figure 4A, *p* = 0.0014). Nascent and asynchronous ase1-7A spindles are also shorter than WT (Figure 4B and 4C). We investigated whether this is due to a lack of kinesin-5 sliding in the ase1- 7A mutant.

**Figure 4.**
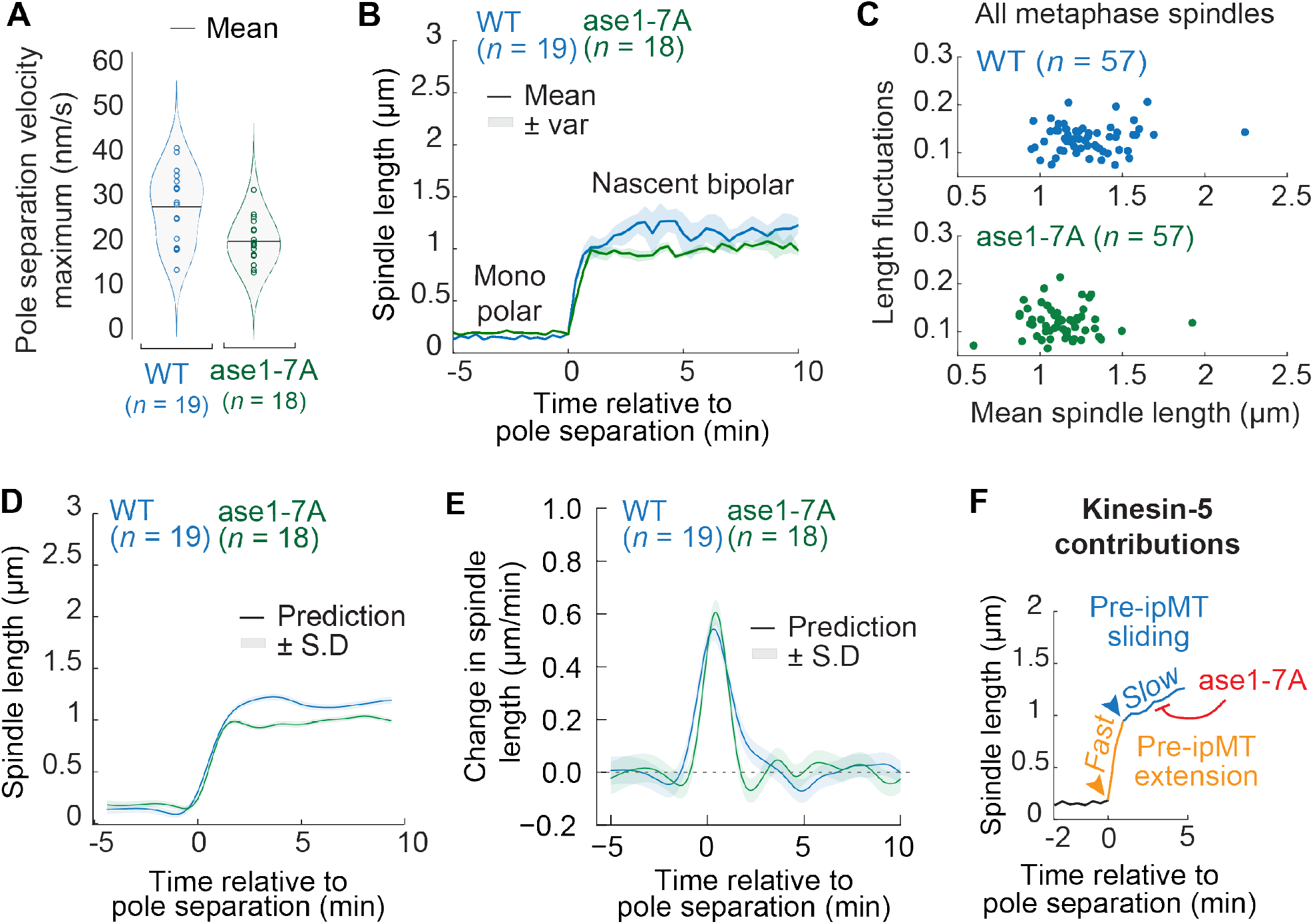
ase1-7A inhibits the kinesin-5 sliding of pre-ipMTs in nascent bipolar spindles (A) Maximum velocity of spindle pole separation during the monopolar-to-bipolar transition in WT and ase1-7A cells. (B) Average and variance of individual WT and ase1-7A trajectories aligned at the monopolar-to-bipolar transition. (C) Spindle length fluctuations as a function of mean spindle length for individual asynchronous WT and ase1-7A cells. (D) Gaussian process regression of spindle length data. The spindle length as a function of time after spindle pole separation was estimated using a gaussian process (with a Matern kernel) for both wild type (blue) and ase1-7A (green) cells. Shaded region shows standard deviation of regression at each timepoint. (E) The derivative of spindle length with respect to time after spindle pole separation, computed from the same gaussian process as in (D). Line is mean and shaded region is the standard deviation of the derivative of the gaussian process. (F) Extension of pre-ipMTs crosslinked by kinesin-5 drives the fast elongation of the spindle, while kinesin-5 sliding of pre-ipMTs drives a slower elongation phase. This slow growth is inhibited by ase1-7A.

We have previously shown that crosslinking alone is not sufficient to maintain net spindle elongation and stability following bipolar spindle formation^2^. This requires kinesin-5 sliding. Indeed, spindles in which kinesin-5 sliding was greatly reduced are shorter and more unstable^2^.

Using a gaussian process regression^37^ of the spindle length before, during, and after the monopolar-to-bipolar transition, we found that in WT cells, this transition is a two-component system, with an initial fast extension and a subsequent slower elongation phase (Figure 4D). This is not the case in the ase1-7A mutant, however, in which the slower elongation phase is missing (Figure 4D). Indeed, we found that in the first few minutes following the transition, the derivative of the regression is much higher for the WT spindles than for the ase1-7A spindles (Figure 4E). This suggests that the kinesin-5 sliding of the pre-ipMTs is inhibited in the ase1-7A mutant, resulting in shorter spindles that do not elongate (Figure 4F).

### ase1-7A restores nascent spindle stability in the γtub-Y445D mutant

Because Ase1 is required for the viability of the γtub-Y445D mutant, we reasoned that it may compensate for the mis-localization of Cin8. We therefore investigated if the ase1- 7A mutant can restore stability to γtub-Y445D spindles. The velocities of the transition in WT, γtub-Y445D, and ase1-7A; γtub-Y445D cells are in agreement with each other (Figure 5A, *p* = 0.2676), consistent with kinesin-5 crosslinking being intact in the γtub- Y445D mutant. We indeed found that inhibiting Ase1 phosphorylation partially rescues the nascent spindle collapses observed in γtub-Y445D cells (Figure 5B). Nascent ase1- 7A; γtub-Y445D spindles are significantly more stable than γtub-Y445D nascent spindles (Figure 5C).

**Figure 5.**
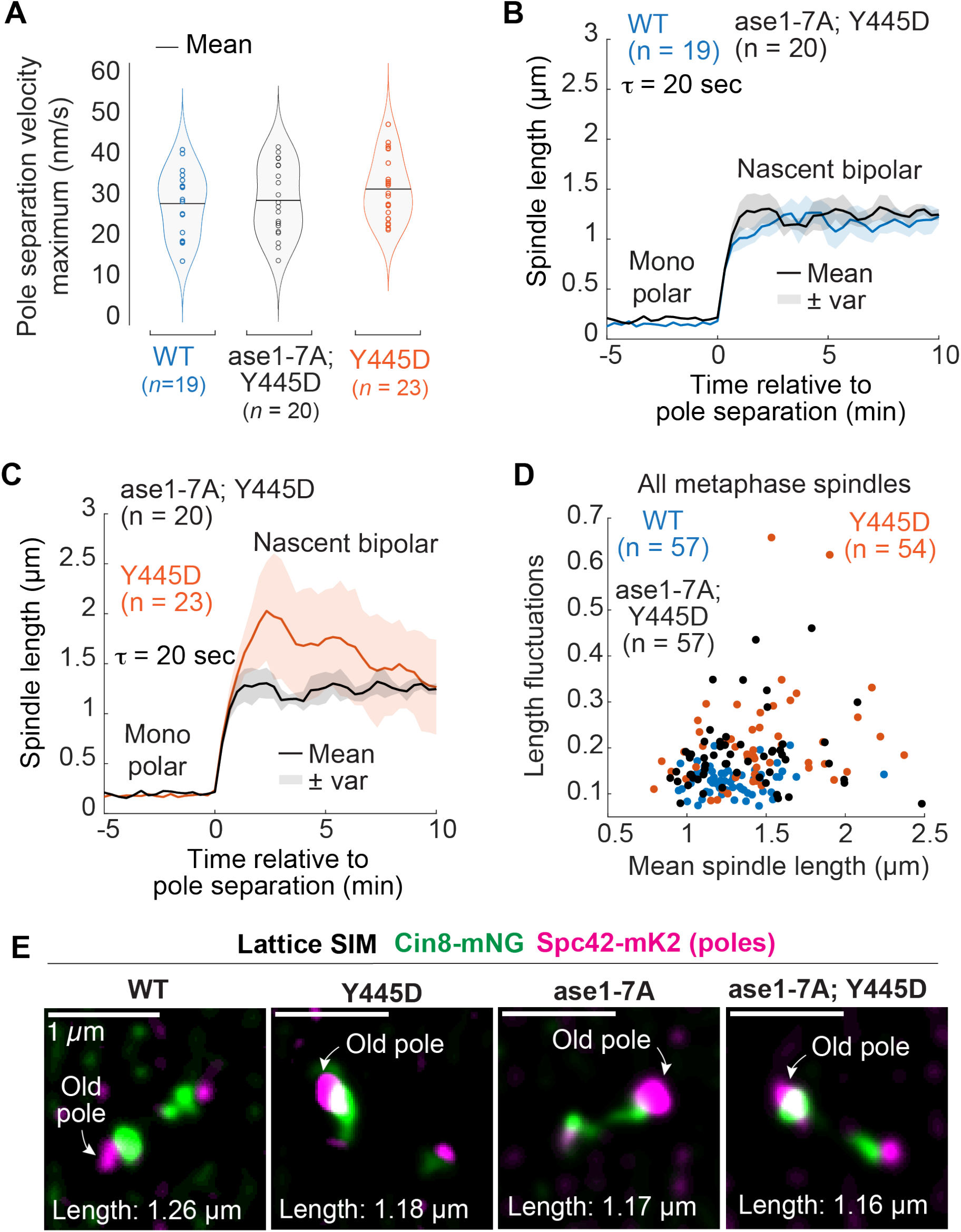
Early recruitment of Ase1 restores nascent spindle stability in **γ**tub-Y445D cells (A) Maximum velocity of spindle pole separation during the monopolar-to-bipolar transition in WT, γtub-Y445D and ase1-7A; γtub-Y445D cells. (B) Average and variance of individual WT and ase1-7A; γtub-Y445D cell trajectories aligned at the monopolar-to- bipolar transition. (C) Average and variance of individual γtub-Y445D and ase1-7A; γtub-Y445D cell trajectories aligned at the monopolar-to-bipolar transition. (D) Spindle length fluctuations as a function of mean spindle length for individual asynchronous WT, γtub-Y445D, and ase1-7A; γtub-Y445D cells. (E) Lattice SIM images of Cin8-mNG on WT, γtub-Y445D, ase1-7A, and ase1-7A; γtub-Y445D cells.

However, we found that asynchronous ase1-7A; γtub-Y445D pre-anaphase spindles are significantly more unstable than WT spindles (Figure 5D, *p* = 2×10^-6^) and have a stability consistent with that of γtub-Y445D pre-anaphase spindles (Figure 5D, *p* = 0.1147). This suggests an inability to extend the short antiparallel overlaps present in nascent spindles through kinesin-5 sliding. Together, these results suggest that Ase1 is capable of compensating for kinesin-5 function early in spindle formation but is incapable of compensating for kinesin-5 sliding, given that it is not a motor protein.

Furthermore, using lattice SIM, we found that as in γtub-Y445D cells, Cin8-mNG localization is biased to the old pole on ase1-7A; γtub-Y445D spindles (Figure 4E). This suggests that the increase in stability observed in nascent ase1-7A; γtub-Y445D spindles is due to the early recruitment of Ase1 and not to Ase1-mediated recruitment of Cin8.

### Changes in spindle stability in relation to centromere attachment reveal the rate of midzone assembly

The midzone stabilizes the spindle and offsets the inward tension generated by the kinetochores. Its formation is required for proper chromosome attachment. However, the rate of formation of the midzone has not been measured.

Nascent, early, and late metaphase spindles can be defined based on their distinct average length regimes. Metaphase spindles are significantly longer than nascent spindles, with an increase in length from 1.1 ± 0.2 µm to 1.4 ± 0.1 µm (*n* = 20, *p* = 3×10^- 7^, Figure 6A and 6C). We found that the time between bipolar spindle formation and anaphase onset varies between 12 and 40 minutes (Fig 6B). Furthermore, nascent spindles (monopolar-to-bipolar transition + 10 minutes) did not transition to anaphase, suggesting that biorientation is inhibited by the lack of a stable midzone during this period.

**Figure 6.**
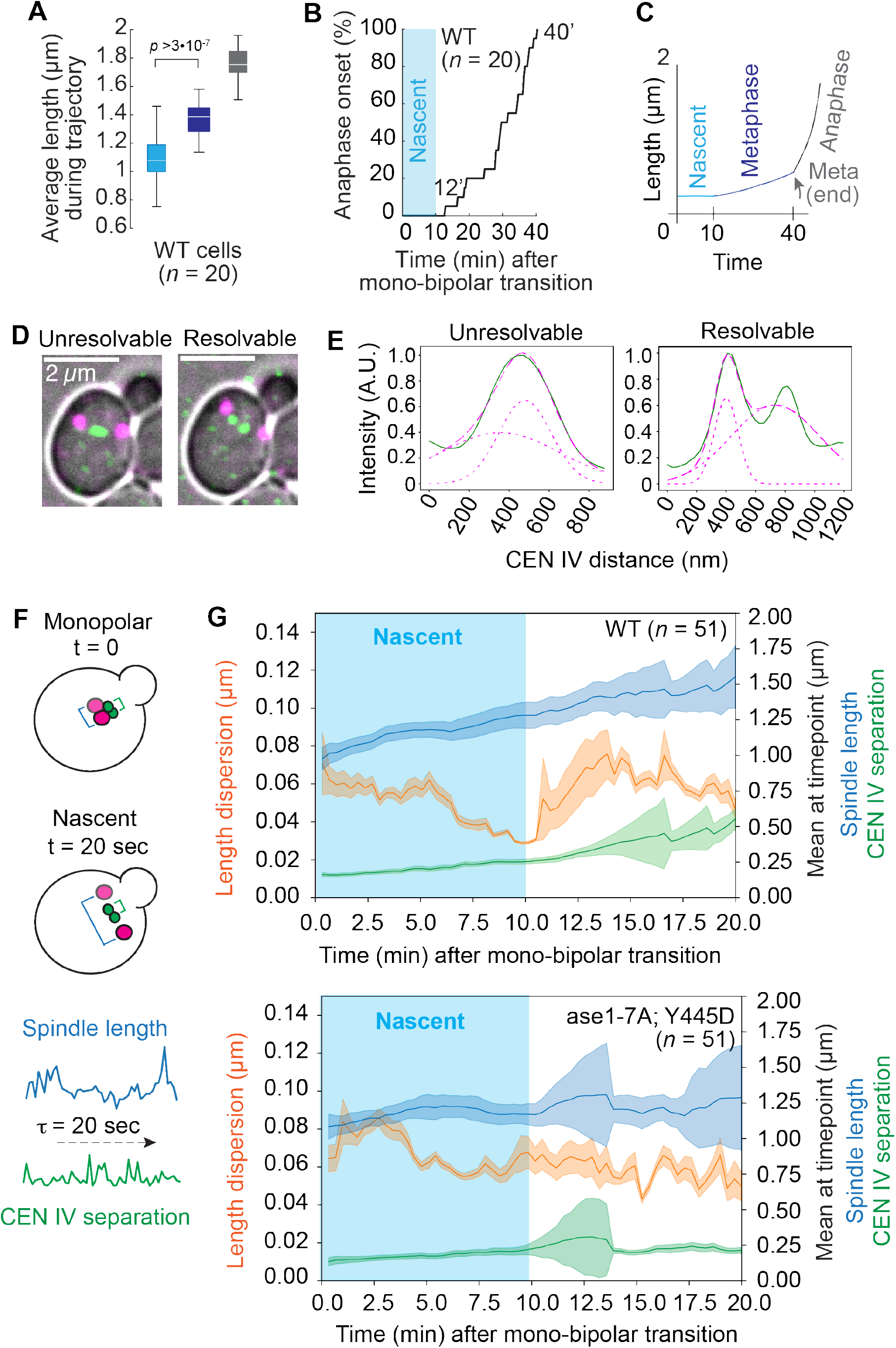
Changes in spindle stability in relation to centromere attachment reveal the rate of midzone assembly and its role in spindle function (A) Average spindle length regimes for nascent, metaphase, and late-metaphase spindles. (B) Cumulative distribution of time between bipolar spindle formation and the metaphase to anaphase transition. (C) Average spindle length throughout metaphase at different stages of spindle assembly for WT cells. (D) Representative examples showing resolvable and unresolvable centromere separation. (E) Examples of gaussian mixture model for determining the centromere separation. Normalized intensity value as a function of distance is shown in green with the fitted mixture model shown in magenta. The short dashes show the individual components while the long dashes show the sum of the components. (F) Graphical summary and timeseries representation showing spindle length and centromere separation measurements. (G) Spindle index of dispersion and CEN4 separation correlated with average spindle length since bipolar spindle formation for WT and ase1-7A; γtub-Y445D cells.

To measure the rate of spindle assembly, we correlated spindle fluctuations, centromere separation (Figure 6D and 6E), and spindle length (Figure 6F) with time elapsed since bipolar spindle formation. Length dispersion (variance of spindle length divided by the mean spindle length) was used to measure the magnitude of spindle fluctuations and infer the formation of the spindle midzone. Centromere separation was used as a proxy for the tension applied to the kinetochores and the beginning of chromosome attachment.

We found, as expected, that nascent spindles are unstable (Figure 6G). For the first ten minutes following bipolar spindle formation, spindle dispersion decreases. We associate this stabilization with the formation of the midzone preceding the generation of inward tension by chromosome attachment. About ten minutes after bipolar spindle formation, we observed a marked increase in spindle dispersion, correlated with an increase in centromere separation (Figure 6G). We attribute these spindle fluctuations to the generation of forces associated with the start of the process of error-prone chromosome attachment.

### ase1-7A cannot stabilize spindles during centromere attachment

We found that the stability of nascent spindles is increased in the ase1-7A; γtub-Y445D mutant relative to the γtub-Y445D mutant (Figure 5C). This suggests that Ase1 is capable of compensating for a defect in kinesin-5 function in early spindle assembly. To determine if this mutant is also capable of converting the pre-ipMTs into ipMTs, we asked if these spindles remained stable during sister chromatid biorientation.

We found that in the ase1-7A; γtub-Y445D mutant, the spindle stabilization observed at approximately ten minutes following bipolar spindle formation in WT spindles does not occur (Figure 6G). Instead, while there is a slight decrease in spindle dispersion a few minutes after bipolar spindle formation, the spindle dispersion remains approximately constant after this time. Furthermore, approximately 12 minutes after bipolar spindle formation, the centromere signal collapses into a single focus. This decrease in the distance between the centromere markers is not observed in WT spindles (Figure 6G). This suggests that the ase1-7A; γtub-Y445D mutant is capable of early spindle assembly but cannot form a stable midzone capable of withstanding the inward tension generated by chromosome attachment. This is possibly due to a lack of kinesin-5 sliding activity in the ase1-7A; γtub-Y445D mutant. This implies that ase1-7A; γtub-Y445D spindles are unable to convert the pre-ipMTs into ipMTs at a similar rate as WT spindles, which causes this observed instability.

## DISCUSSION

### **γ**-Tubulin phosphoregulation may be a mechanism for tuning the number of ipMTs that are formed

Surprisingly, despite γ-tubulin being located at the minus ends of MTs, our results strongly suggest that the formation of ipMTs by Cin8 at the plus ends of MTs is reduced in the γtub-Y445D mutant. The γtub-Y445D mutation produces phenotypes consistent with a defect in Cin8 function; the stability of bipolar spindles is reduced relative to WT cells, and the MT crosslinking protein Ase1 which is involved in forming the ipMTs and is essential in the absence of Cin8 is essential in γtub-Y445D cells.

The γ-tubulin Y445 residue is known to be phosphorylated during spindle assembly^13,14^. Given the evolutionary conservation of this site, it is likely that its regulation procures an advantage over evolutionary timescales.

The detrimental effect of the γtub-Y445D mutation on fitness suggests that phosphorylation of Y445 is a rare event and that it may only occur in a subset of γ-tubulin molecules. It may be a mechanism that allows the number of ipMTs to be limited, allowing free microtubules to search for and attach to kinetochores. Such mechanisms to ensure tight control of ipMT number would be advantageous to the cell, given the importance of accurate chromosome segregation.

### The conformation of the **γ**CT may control kinesin-5 activity

Previous work has shown that the γtub-Y445D mutation drives the γCT to sample unique extended conformations that are not observed in WT controls^16^. We hypothesize that the phosphorylation state of Y445, which alters the conformation of the γCT, controls kinesin- 5 interactions with MTs through an unknown mechanism that controls the number of ipMTs that will be formed. This mechanism could be direct or indirect.

One possibility is that the extended conformations of the γCT that are present when Y445 is phosphorylated selectively recruit Cin8 to the MTs or inhibit its release from the MT minus ends. Cin8 would be sequestered at the spindle poles in γtub-Y445D cells and depleted from the spindle midzone. Antiparallel MT pairing and the formation of the ipMTs would be inhibited, resulting in spindle instability. γCT phosphorylation could provide additional control over the number of MTs that will pair and form the ipMTs as the spindle forms. The γCT of human γ-tubulin is shorter than that of budding yeast, ending shortly after Y443 (Y445 in budding yeast). It is possible that motor proteins such as Cin8 have evolved interactions with this extended tail when they walk to the minus ends of the MTs. Cut7, the fission yeast kinesin-5, binds to γ-tubulin, and this binding contributes to its spindle pole localization^38^.

We also speculate that phosphorylation of Y445, which is expected to increase the radius of gyration of the γCT, could act as a scaffold to regulate the exchange of proteins that influence MT organization and/or dynamic instability. A model for the centrosome as a biomolecular condensate that promotes MT nucleation and specification has previously been proposed^39^. As well, the recruitment of Kip2 at the minus ends of MTs in budding yeast has been linked to the control of MT length, and thus to events at the plus ends of MTs^40^.

### Ase1 crosslinking of pre-ipMTs inhibits kinesin-5 sliding in nascent spindles

We have shown that both nascent and asynchronous ase1-7A spindles are shorter than WT (Fig. 4B and 4C). Furthermore, the velocity of the monopolar-to-bipolar transition is reduced in this mutant (Fig. 6A). This may be due to competition between kinesin-5 and Ase1, with Ase1 opposing the sliding forces generated by kinesin-5. It also suggests that the ase1-7A rescues the stability of nascent γtub4-Y445D spindles not through Ase1- mediated recruitment of Cin8^36^, but through the crosslinking capabilities of Ase1 itself.

Spindle length is stereotyped and calibrated to cell volume^41^. An inability to extend spindle length through kinesin-5 sliding may have consequences for the cell. For example, some of the chromosome arms in a short spindle may not fully leave the spindle midzone during anaphase, with lagging chromosomes that could result in aneuploidy.

### Spindle stability can be used as a tool to understand midzone formation

We report that spindles are stabilized approximately 10 minutes following bipolar spindle formation, from which can infer the formation of the ipMTs (Fig. 6G). Furthermore, we find that the shortest time between bipolar spindle formation and anaphase is about 12 minutes (Fig. 6B). This suggests that until the spindle has formed an adequate midzone, biorientation is inhibited. It also suggests that as soon as the midzone forms, chromosome attachment is underway, and some spindles are ready to undergo anaphase.

We also find that in ase1-7A; γtub4-Y445D spindles, the distance between the centromere markers increases and subsequently collapses in a very concerted manner (Fig. 6G). This suggests that this mutant has a defect in tension generation rather than in chromosome attachment. Furthermore, this centromere separation and collapse begins 10 minutes after bipolar spindle formation, which is a few minutes before centromere separation starts to be observed in WT cells (Fig. 6G). This could indicate that chromosome attachment is more efficient when the search space for forming attachments is reduced. Thus, spindle length at the end of metaphase may be calibrated for subsequent events in anaphase.

## ACKNOWLEDGEMENTS

The authors thank members of the Vogel lab for discussions and their comments on the manuscript. We thank Katherine Morelli for her contribution of the multi-gaussian fitting method and Angela Zhao and Elisa Ammon for valuable scientific discussions. S.S. is supported by a fellowship from the Fonds de recherche du Québec, Nature et Technologies (FRQNT, award #313160) and the Lorne Trottier Science Accelerator Fellowship. K.A. is supported by a scholarship from Qatar University. This research was supported by a grant from the Canadian Institutes of Health Research PJT-1666078 to J.V.

## AUTHOR CONTRIBUTIONS

Conceptualization, J.V. and S.S.; Strain construction, S.S., K.A., Z.E.; Data analysis, S.S., S.M., K.A.; Live-cell microscopy, S.S., S.M., K.A.; Writing - Original Draft, S.S. and J.V.; Writing – Review & Editing, S.S., J.V., S.M., K.A.; Supervision, J.V.; Funding Acquisition, J.V.

## METHODS

### KEY RESOURCES TABLE

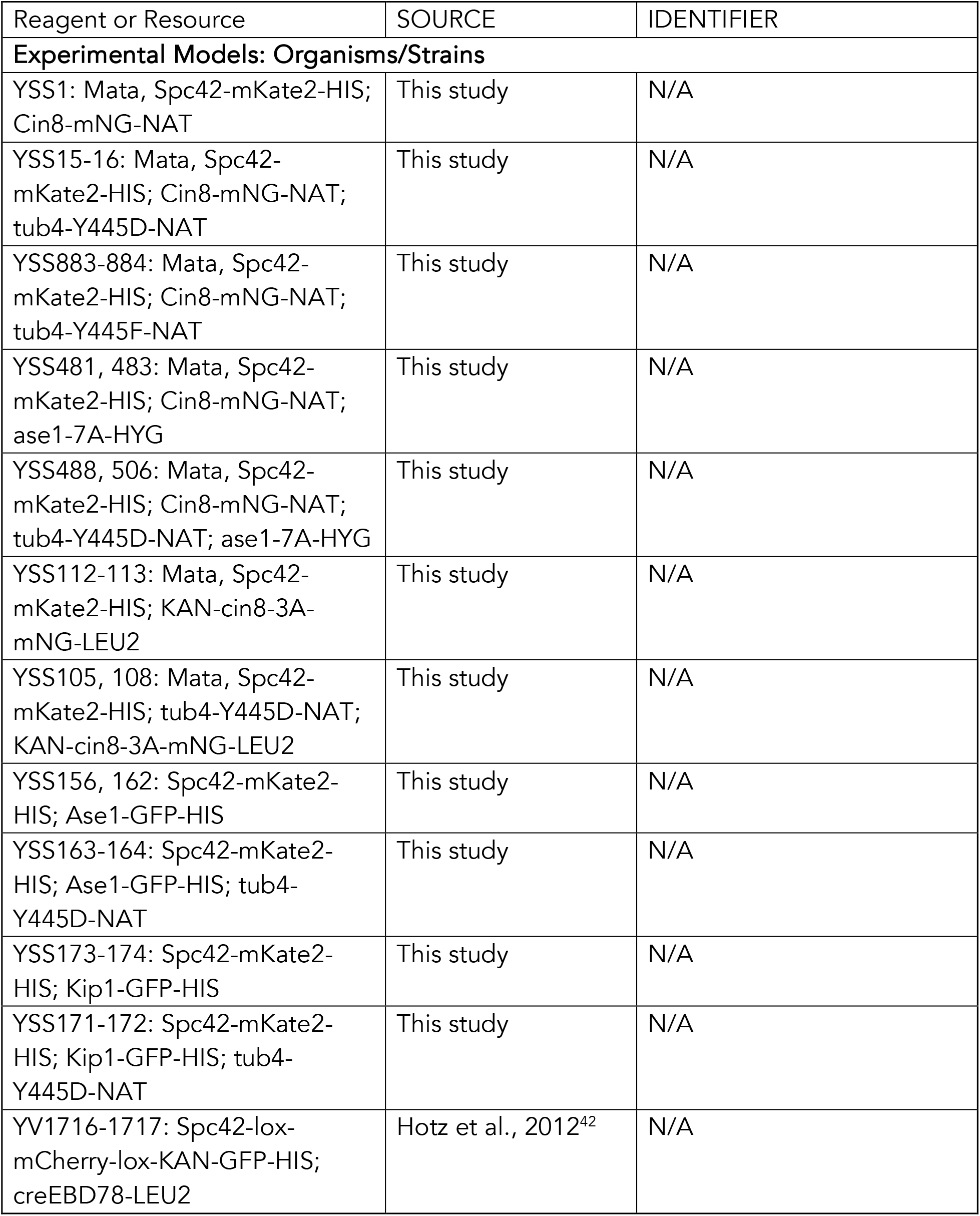
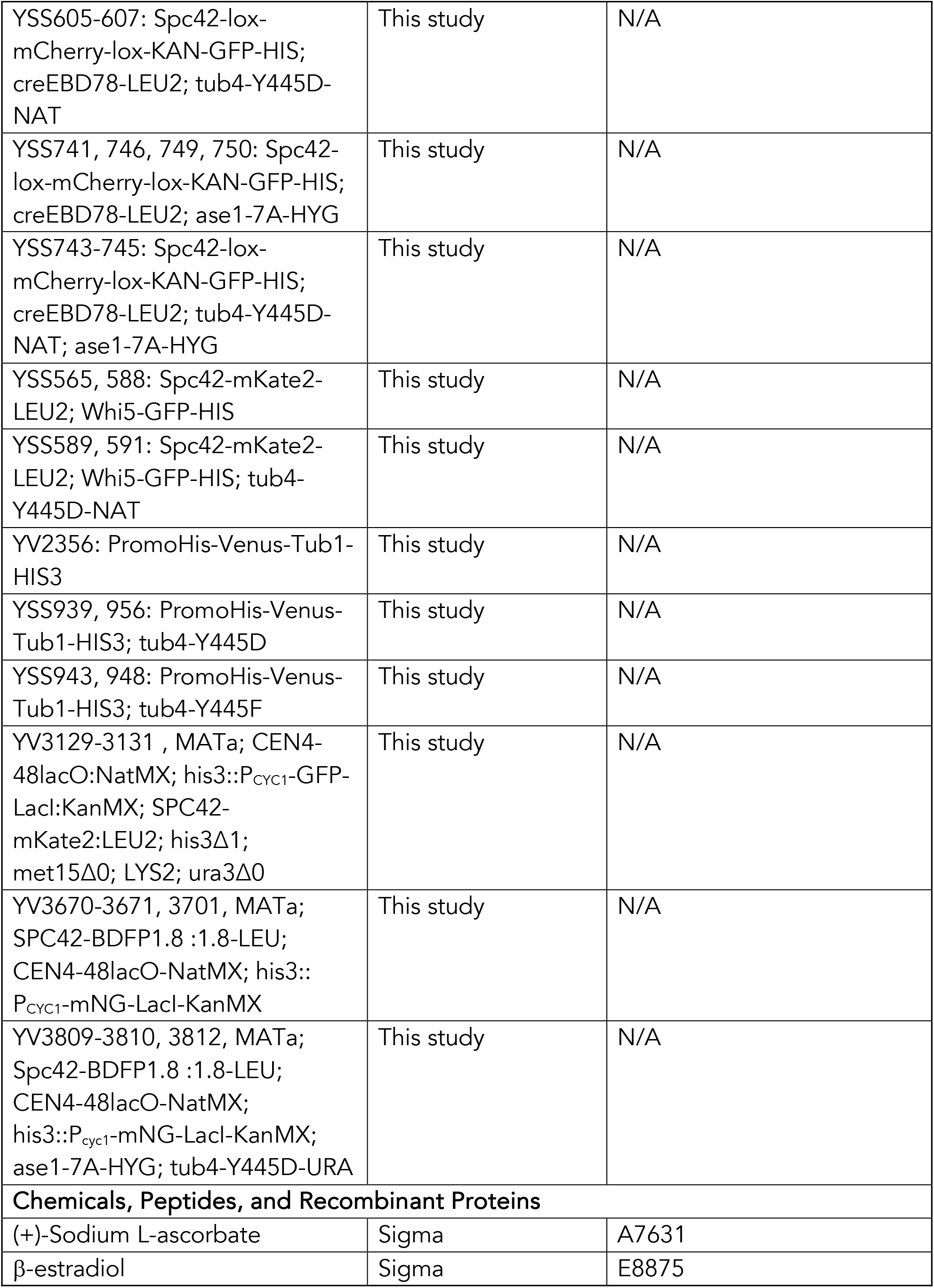
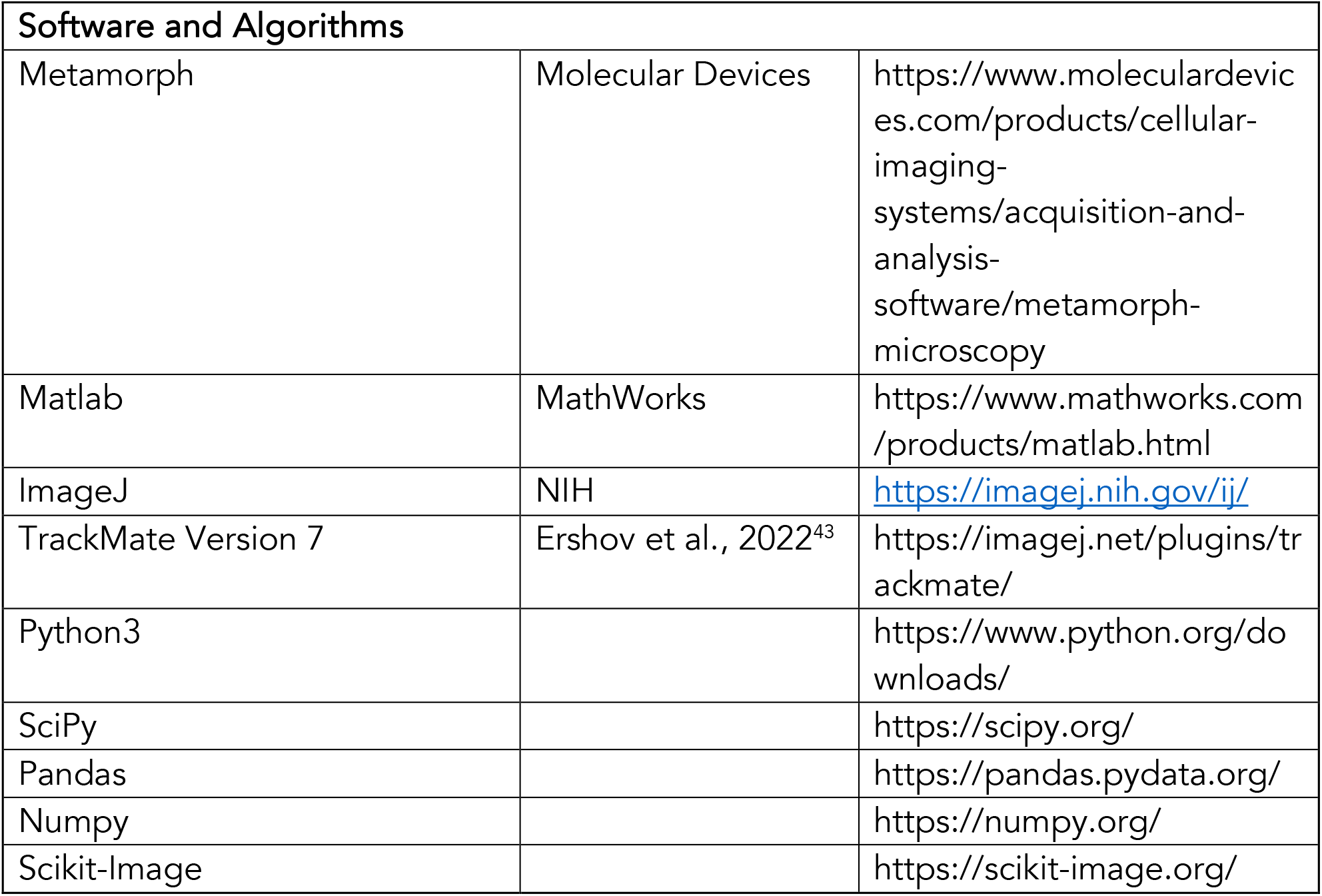

### LEAD CONTACT AND MATERIALS AVAILABILITY

Lead contact and materials availability Further information and requests for resources and reagents should be directed to and will be fulfilled by the lead contact, Jackie Vogel.

### EXPERIMENTAL MODEL AND SUBJECT DETAILS

#### Yeast strains

All yeast strains in this study are derivatives of BY4741 and are found in the above key resources table.

#### METHOD DETAILS

### Live cell microscopy

Yeast strains were grown at 25°C in 5 mL cultures of synthetic complete (SC) media^44^ overnight, diluted in the morning (OD600 ∼ 0.2) and supplemented with 2mM ascorbic acid and allowed to grow to mid log phase. Low fluorescence SC was used for the super-resolution Cin8-mNG and VFP-α1 tubulin images (as noted below), in which Formedium low-fluorescence yeast nitrogen base is used and autoclaving is replaced by filter sterilization. All live cell imaging was carried out at 25°C. Cells were prepared for imaging as described previously^2,3^.

The confocal imaging of Spc42-mKate2, Cin8-mNG, Kip1-GFP, and Ase1-GFP was carried out on custom-built dual-camera spinning disk confocal microscope. The system consists of the following components: a Leica DMi8 inverted microscope with a CSU10 confocal scanner unit (Yokogawa), a HCX PL APO 100x/1.47 OIL CORR TIRF objective (Leica), two iXon Ultra 512 x 512 EMCCD cameras for simultaneous acquisition (Andor), an ASI three axis motorized stage controller and MCL nano-view piezo stage, 488 nm and 561 nm solid state OPSL lasers linked to a Borealis beam conditioning unit. MetaMorph (Molecular Devices) was used for microscope control and image acquisition. Images were acquired in stream mode, with 30 z-slices at 200 nm intervals forming 5.8 µm z-stacks and an exposure time of 50 ms. The cells were imaged for 10 mins with a 10 s timestep between images (61 timepoints).

For the imaging of initial spindle formation with the Cre-Lox fluorophore-switching system, Cre recombination was induced by adding 1 mM β-estradiol to the SC media 120 mins before imaging to ensure that new GFP spindle poles were present. The microscopy setup described in the previous section was used. Images were acquired in stream mode, with 30 z-slices at 200 nm intervals forming 5.8 µm z-stacks and an exposure time of 50 ms. The cells were imaged for 20 mins with a 20 s timestep between images (61 timepoints).

The lattice SIM VFP-α1 tubulin and lattice SIM Cin8-mNGr and Spc42-mKate2 images were acquired on an Elyra 7.2 dual-camera lattice SIM microscope (Zeiss) with the following components: a 100x oil objective (alpha Plan-Apochromat 100x/1.57 Oil-HI DIC Korr M27 Elyra), two PCO.edge sCMOS version 4.2 CLHS cameras (Excelitas), a Z piezo stage, and solid state 488 nm and 561 nm lasers linked to an LCS-BU liquid cooling system. Sapphire coverslips (Zeiss) were used. The software used for microscope control and image acquisition was ZEN 3.0 SR Black (Zeiss). 13 lattice SIM phases were used for the acquisitions. Each 3.48 µm z-stack was composed of 41 z-slices taken at 87 nm intervals and an exposure time of 2 ms for the VFP-α1 tubulin images and 5 ms for the Cin8-mNG images. Images were processed using the SIM^2^ algorithm and weak live processing, which is chosen based on the signal-to-noise ratio.

To measure the timing of bipolar spindle formation relative to START, cells were deposited on a 1% agarose SC pad contained in a gene frame (ThermoScientific). The microscope used for acquisitions was an Olympus IX83 microscope with the following components: a 100x oil objective lens (Olympus Plan Apo 100x NA 1.40 oil), an Orca-Flash 4.0 sCMOS camera (Hamamatsu), a NanoScanZ piezo (Prior Scientific), and an X-Cite 120 LED lamp. The mCherry-WF and FITC-WF filters were used. cellSens Dimension (Olympus) was used for microscope control and image acquisition. 11 z-slices at 0.5 µm intervals were acquired to form 5 µm z-stacks with an exposure time of 100 ms. 120 stacks were acquired at 2 min intervals for a total imaging time of 4 hrs.

### Time spent in metaphase

For the measurement of the time spent in metaphase, 6 μL of the concentrated culture was plated on a synthetic complete 1% agar Gene Frame. To increase the signal-to-noise ratio of the data collected, the coverslip (Ted Pella 0.16-0.19mm) was cleaned with acetone/methanol/water washes, and remaining contaminants were removed from the glass using a plasma oven (PE-50LF-XL Plasma Etch). The acquisitions were carried out on the same Elyra 7.2 microscope described above with 0.2 µm intervals for 13 slices with an exposure time of 25 ms, with images taken at 20 s intervals for 1 hr.

### Centromere and chromosome attachment

For the data used to calculate the index of dispersion, 10 μL of the concentration culture was plated on a 24mm Deckgläser coverslip rested on a custom circular ring (courtesy of the Barral lab) then covered with a layer of SC agar. The setup was then sealed with a Deckgläser coverslip and images were acquired on an Elyra 7.2 microscope as described above with 0.2 µm intervals for 13 slices with an exposure time of 25 ms, with images taken at 20 s intervals for 20 mins.

## QUANTIFICATION AND STATISTICAL ANALYSIS

### Tracking and analysis of the spindle length and stability of asynchronous bipolar spindles

The positions of the spindle poles in three dimensions were computed using TrackMate (FIJI)^43^. Only cells for which the spindle poles were tracked for at least 45 out of the 61 timepoints were retained. The brighter Spc42-mKate2 foci was identified as the old spindle pole^3^. The spindle fluctuations were computed as the standard deviation of the spindle lengths for individual cells.

### 3D Kymograph Generation

Kymographs were generated using the positions of the spindle pole bodies (generated using TrackMate (FIJI)^43^) and raw image data. For each 3D frame of each video, comprising of an X, Y, and Z dimensions, the position of the spindle poles is computed with respect to the image and a single column of a kymograph is generated. The location of the spindle pole bodies is measured in microns, so the coordinates were divided by the voxel size to represent the coordinates in terms of the pixels in the image. Once the spindle pole positions were identified in pixel coordinates, the values of the pixels in a single pixel wide straight-line path from the center of the old spindle pole body to the center of the new spindle pole body were recorded. Once an array of pixel values was generated for each 3D frame of a video, the arrays were expanded to a uniform length using a linear interpolation and the columns were arranged sequentially to form a kymograph.

Once a kymograph was generated to each cell in a given condition, they were merged into a single kymograph. Then, each kymograph was resized into a 2D array of 256 by 256 pixels (SciKit-Image). This value was chosen because each dimension is significantly larger than any individual kymograph, so no information would be lost. Image resizing was done using the “nearest neighbor” method to determine new pixel values. Finally, the corresponding values in each individual kymograph were averaged to create an average kymograph for each condition.

### Anlaysis of the time interval between START and bipolar spindle formation using Whi5-GFP and Spc42-mKate2

Completion of the export of Whi5-GFP from the nucleus in G1 was used to identify START. Z-stacks of the Whi5-GFP channel were sum projected in FIJI. START was identified by visual inspection as the first timepoint when the nuclear Whi5-GFP signal was indistinguishable from the cytoplasmic Whi5-GFP signal. Spc42-mKate2 was used to visualize the spindle poles and bipolar spindle formation was determined to be at the first time point when two distinct Spc42-mKate2 foci could be resolved by visual inspection in maximum projection images (FIJI). The time elapsed between these two events was determined to be the time interval between START and bipolar spindle formation.

### Analysis of initial spindle formation

The positions of the spindle poles in three dimensions were computed using TrackMate (FIJI)^43^. The onset of the monopolar-to-bipolar transition was determined by visual inspection. Only cells with trajectories lasting at least 10 minutes following bipolar spindle formation were retained. The maximum velocity during the transition was determined to be the maximum velocity between two timepoints in the trajectory.

### Spindle length computation

The spindle length was computed by taking the Euclidian distance between the coordinates of the old and new spindle pole. These coordinates were obtained using TrackMate (FIJI)^43^.

### Centromere separation computation

The pixel intensity values of the microscope image were recorded by drawing a 10 pixel wide line over the image in Fiji and recording the intensity along that line. If there are two resolvable foci, then the line was drawn through both of their centers. If there was only a single resolvable focus, then the line was drawn over the center of the focus in the direction of maximal stretching, determined by inspection. If there is a single focus with no discernable longest axis, then the line was drawn through the center of the focus in the direction parallel to the mitotic spindle.

Then, for each timepoint a gaussian mixture model with two components was fit to the data with the constraint that one component could be no less than 0.3 times the size of the other and that they could not have a variance larger than 300 nm. The mixture model was fit using the dual annealing algorithm (Python library SciPy) to minimize the mean squared error. Finally, the distance between the means of the two components was calculated to obtain the centromere separation along the line drawn in the image. This was done for each timepoint in each cell where the centromere was visible.

### Mean spindle length, spindle length dispersion, and mean centromere separation calculation

For each cell at each timepoint, the instantaneous spindle length and centromere separation were computed (see above) resulting in two time series for each cell. Each statistic at each timepoint was computed using a three-minute window centered on the timepoint. The mean and variance were computed over this window (using the Python libraries NumPy and pandas) and the dispersion was computed by dividing the variance by the mean.

### Spindle length gaussian process regression and derivatives

The spindle length as a function of time since the monopolar-bipolar transition was estimated using a gaussian process with a Matern kernel^37^. The bounds for the hyperparameters were selected to be [10^-4^, 10^4^] for the amplitude parameter, [10^-4^, 10] for the stiffness hyperparameter, and [10^-2^, 10^2^] for the measurement error hyperparameter. The gaussian process was fit to the dataset of all cell trajectories at one time and mean and standard deviation of the spindle length and spindle length derivative were computed for all timepoints using previously developed methods^37^.

---

**Figure S1.**
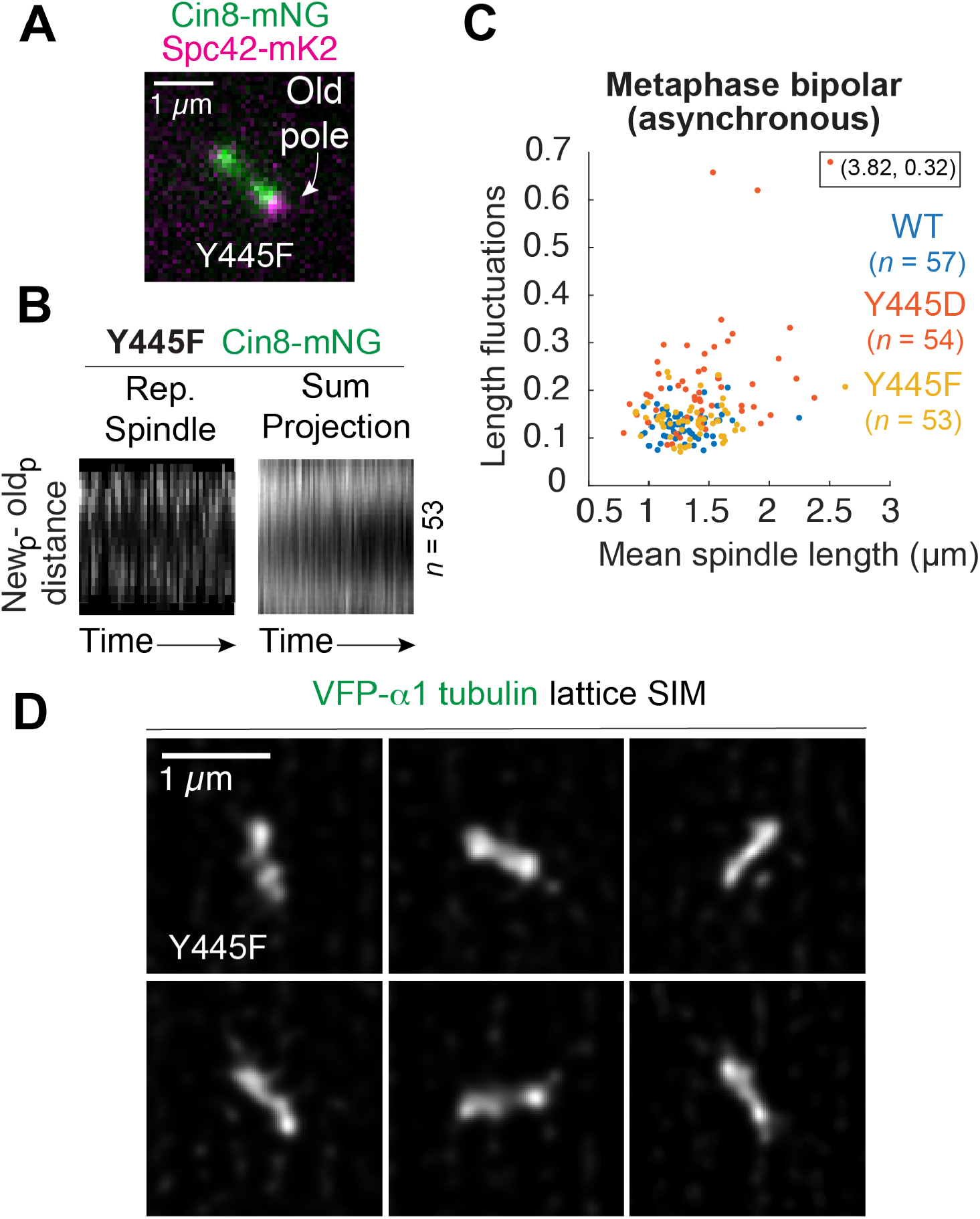
The **γ**tub-Y445F mutation does not cause a bias of Cin8-mNG to the old pole, spindle instability or disrupt ipMT formation (A) Distribution of Cin8-mNG on a γtub-Y445F spindle. (B) Dynamic localization of Cin8-mNG during metaphase in γtub-Y445F cells. Kymographs for a single representative spindle and a summed projection are shown. (C) Spindle length fluctuations as a function of mean spindle length for individual asynchronous γtub-Y445F cells. (D) Lattice SIM analysis of midzone structure in γtub-Y445F cells.

**Figure S2.**
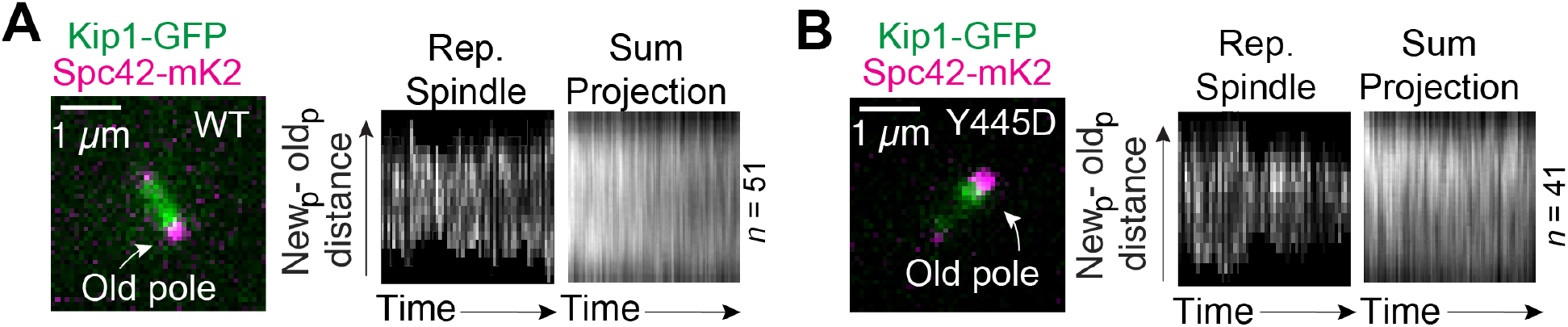
Kip1-GFP localization is biased to the old pole in **γ**tub-Y445D cells (A)-(B) Dynamic localization of Kip1-GFP during metaphase in WT and γtub-Y445D cells. Kymographs for a single representative spindle and a summed projection are shown for each condition.

**Figure S3.**
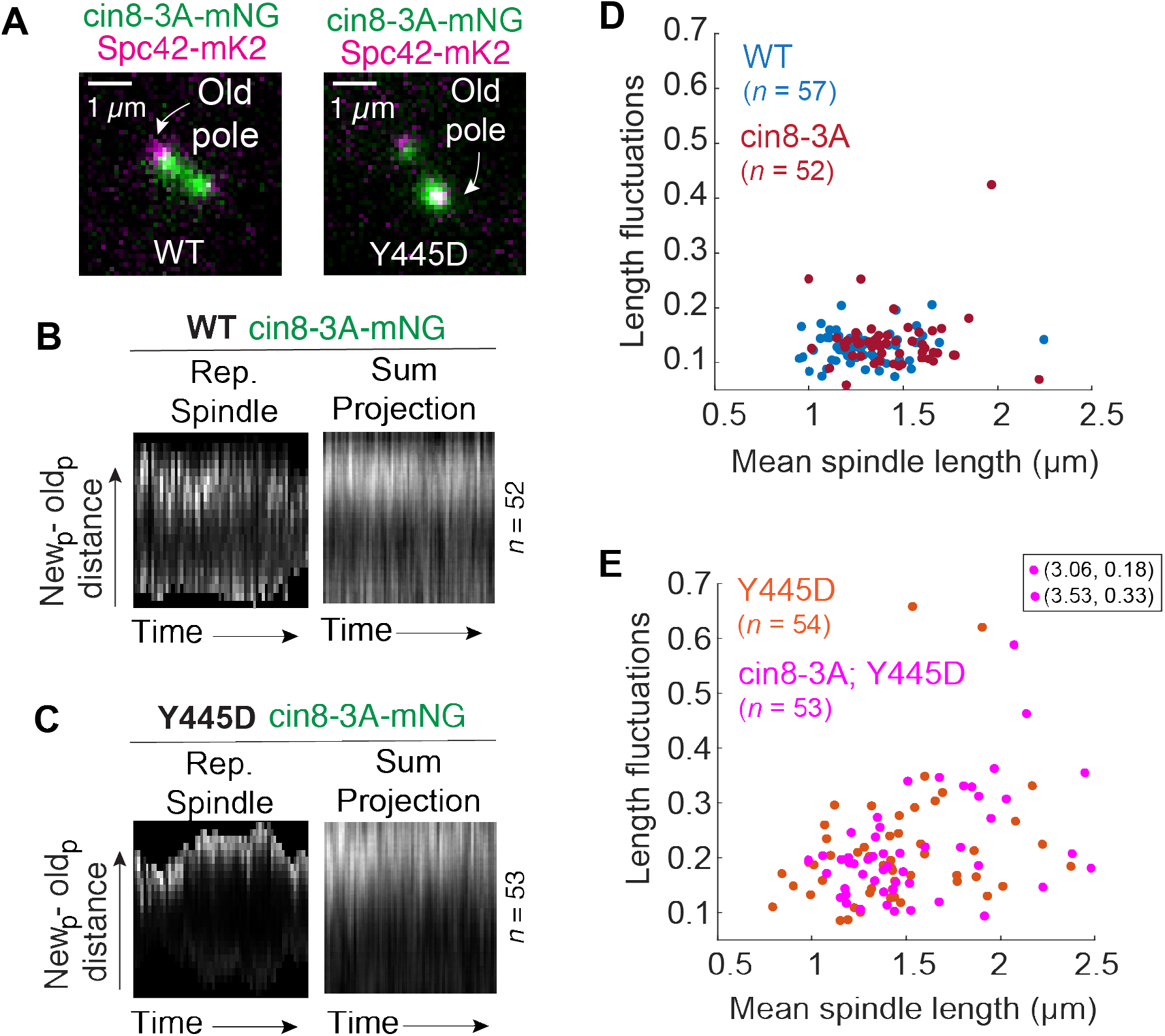
The bias of Cin8-mNG to the old pole is increased in cin8-3A; **γ**tub-Y445D cells (A) Distribution of cin8-3A-mNG on cin8-3A and cin8-3A; γtub-Y445D spindles. (B)-(C) Dynamic localization of Cin8-mNG during metaphase in cin8-3A and cin8-3A; γtub-Y445D cells. Kymographs for a single representative spindle and a summed projection are shown for each condition. (D)-(E) Spindle length fluctuations as a function of mean spindle length for individual asynchronous WT, γtub- Y445D, cin8-3A, and cin8-3A; γtub-Y445D cells.

**Figure S4.**
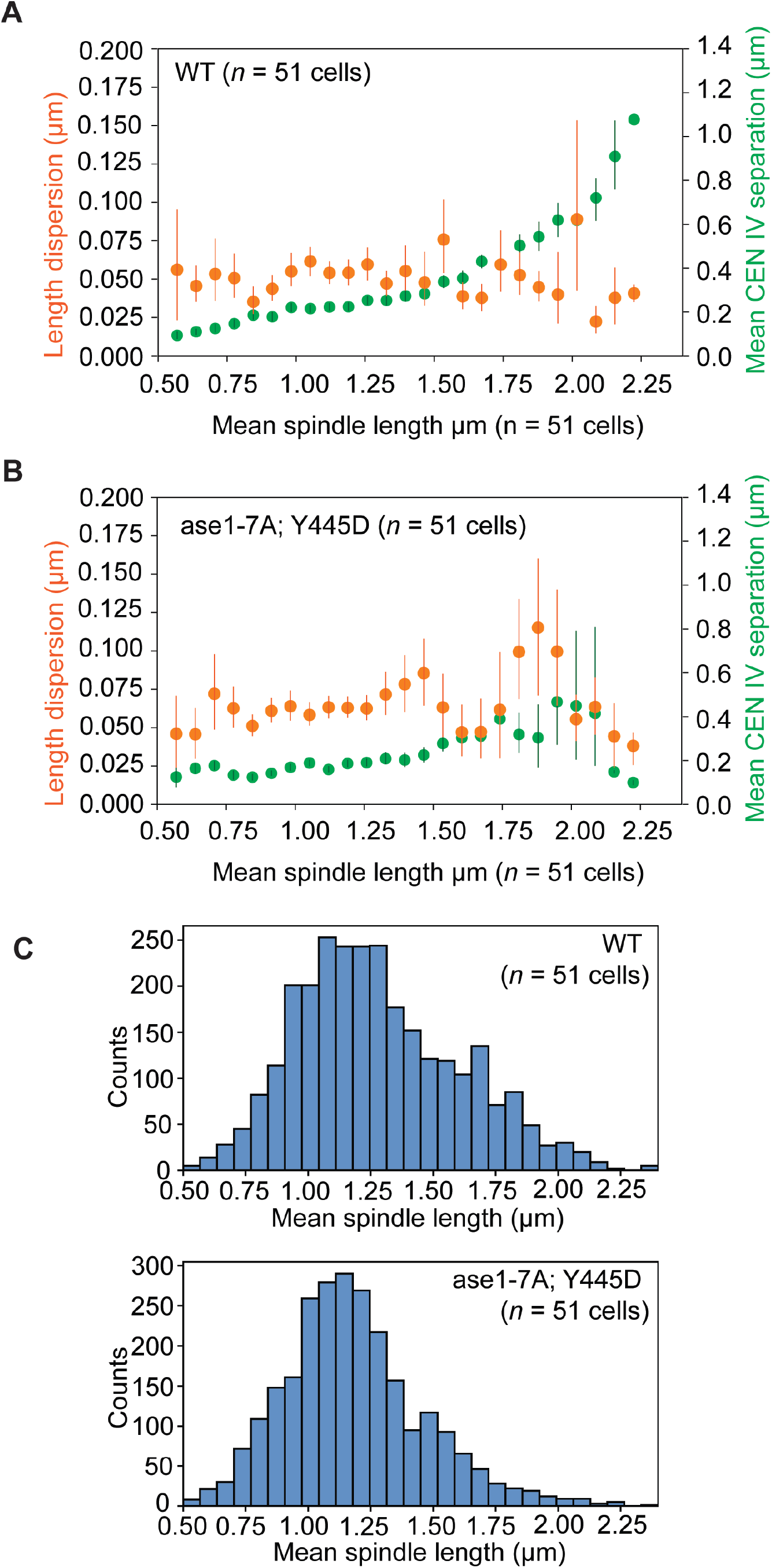
Length dispersion and centromere separation correlated with mean spindle length (A) Spindle index of dispersion correlated with mean CEN4 (mNG-LacI bound to LacO) separation over mean spindle length for WT cells. (B) Spindle index of dispersion correlated with mean CEN4 (mNG-LacI bound to LacO) separation over mean spindle length for ase1-7α; γtub-Y445D cells. (C) Histogram showing spindle length distribution for both WT and ase1-7A; γtub-Y445D cells. The number of bins was determined using Sturge’s rule.

